# A functionally validated TCR-pMHC database for TCR specificity model development

**DOI:** 10.1101/2025.04.28.651095

**Authors:** Marius Messemaker, Bjørn P.Y. Kwee, Živa Moravec, Daniel Álvarez-Salmoral, Jos Urbanus, Sam de Paauw, Jeroen Geerligs, Rhianne Voogd, Ben Morris, Aurélie Guislain, Maike Mußmann, Yaël Winkler, Maxime Steinmetz, Matyas Iras, Eric Marcus, Jonas Teuwen, Anastassis Perrakis, Roderick L. Beijersbergen, Wouter Scheper, Ton N. Schumacher

**Author notes:** These authors contributed equally (Bjørn P.Y. Kwee, Živa Moravec). These authors contributed equally (Wouter Scheper, Ton N. Schumacher).

## Abstract

Accurate prediction of TCR specificity forms a holy grail in immunology and large language models and computational structure predictions provide a path to achieve this. Importantly, current TCR-pMHC prediction models have been trained and evaluated using historical data of unknown quality.

Here, we develop and utilize a high-throughput synthetic platform for TCR assembly and evaluation to assess a large fraction of VDJdb-deposited TCR-pMHC entries using a standardized readout of TCR function. Strikingly, this analysis demonstrates that claimed TCR reactivity is only confirmed for 50% of evaluated entries. Intriguingly, the use of TCRbridge to analyze AlphaFold3 confidence metrics reveals a substantial performance in distinguishing functionally validating and non-validating TCRs even though AlphaFold3 was not trained on this task, demonstrating the utility of the validated VDJdb (TCRvdb) database that we generated. We provide TCRvdb as a resource to the community to support training and evaluation of improved predictive TCR specificity models.

## Introduction

The T cell-based adaptive immune system provides the body with the capacity to eliminate infected or malignant cells. T cells specifically detect antigenic peptides presented by major histocompatibility complex (MHC) molecules through their heterodimeric αβ T cell receptor (TCR). During thymic development, T cells acquire their TCR α and β chains through somatic recombination of V(D)J gene segments, along with random nucleotide insertion and deletion at the resulting gene junctions, generating an estimated 10^8^-10^10^ different TCR clones with distinct reactivity patterns in an individual^1^. This ‘distributed recognition capacity’ equips the human T cell compartment with the capacity to e.g., detect and respond to pathogens, including those yet to emerge, and to eliminate malignant cells that present cancer (neo-)antigens^2^. Ever since the discovery of MHC restriction by Doherty and Zinkernagel^3^ and the cloning of the first T cell receptors by the groups of Mak and Davis^4–6^ immunologists across scientific fields have aimed to identify the antigens and T cell receptors that drive autoimmune disease^7^ and cancer regression^8^, and that can inform vaccine design^9–12^.

Given the essential role of T cell recognition in a variety of human conditions, and given the inability to experimentally map this vast interaction space, *in silico* models that can predict TCR-peptide-MHC reactivity hold a major promise for our understanding of adaptive T cell immunity. As an example, in immuno-oncology, such predictive models may enable the selective boosting of tumor-reactive T cell responses, or may facilitate patient stratification based on the breadth of the tumor-reactive T cell pool. Beyond cancer, predictive models of TCR specificity could advance our understanding of autoimmune disease by revealing which self-antigens are recognized by autoreactive T cells, or aid in rational vaccine design. Ultimately, robust prediction models of TCR specificity may enable the *in silico* design and iterative optimization of synthetic TCRs that are free from the constraints of the natural TCR pool.

Recent advances in deep-learning-based approaches have resulted in protein structure prediction models such as Alphafold and RoseTTAFold that show unprecedented accuracy^13–15^. The remarkable progress in these protein structure prediction models has relied heavily on the large volume of experimental structural information available in the Protein Data Bank (PDB)^16^, serving as a training data resource for deep learning approaches. Importantly, PDB provides multiple standardized validation measures to assess the quality of deposited structures, such as *R*_free_ and the proportion of Ramachandran outliers^17,18^, and development of this standardized validation process has been associated with an increase in the quality of deposited protein structures over time^18^.

As compared to the prediction of protein structures and protein complexes in other fields, progress in predicting reactive (i.e., inducing TCR signaling) TCR-pMHC pairs has to date been modest^19,20^. Part of the complexity of this task is related to the low (µM range) affinity of TCR-pMHC interactions and the fact that the binding site of TCRs is composed of loops with considerable conformational flexibility. However, we speculated that the quality of datasets that have been used to train and evaluate TCR-pMHC predictors could also form a major factor that has limited progress in this field. The primary training and evaluation^21^ data resource for models of TCR specificity has to date been formed by the VDJdb^22^ and IEDB^23^ databases. Importantly, the TCR-pMHC pairs that make up these databases have been identified in a variety of studies using diverse assays and sample sources. With the goal to increase data quality, the TCR-pMHC pairs that are deposited in these databases are curated using assay information. In addition, VDJdb data contain confidence scores based on presumed indicators of assay quality. However, whether such confidence scores hold similar value as e.g., PDB validation metrics such as *R*_free_ has not been established.

With the aim to provide standardized metrics of TCR reactivity, we here develop a scalable semi-synthetic platform to evaluate antigen recognition. We then utilize this platform to demonstrate that the quality of data used to create TCR-pMHC prediction models is concerningly low. We also demonstrate that the performance of established sequence-and structure-based models of TCR specificity has been underestimated because of this issue. Finally, we provide a validated dataset that should allow the more effective training of novel TCR-pMHC prediction models.

## Results

### Exploration of data quality in VDJdb

The VDJdb MHC-I database contains approximately 6,300 TCRαβ entries that are dominated by a modest number of well-studied viral epitopes, with ten epitopes making up about half of the database (Supplementary Table 1). To assess the quality of TCR-pMHC pairs contained in this database, we focused on the HLA-A*02:01-restricted GLCTLVAML (GLC) Epstein–Barr virus (EBV) and YLQPRTFLL (YLQ) severe acute respiratory syndrome coronavirus 2 (SARS-Cov2) epitopes, representing epitopes for which TCR discovery has either been pursued for decades or is restricted to the past few years. For each epitope, a set of TCRs with more similar or distinct sequences was selected. Following manual synthesis and arrayed introduction into CD8^+^ Jurkat T cells, capacity of these TCRs to recognize either the cognate GLC or YLQ epitope was evaluated in co-cultures using CD69 expression as a measure of TCR signaling. Strikingly, of the 49 YLQ-annotated and 41 GLC-annotated TCRs evaluated, the claimed reactivity was only observed in 34.7% and 63.4% of cases, respectively (Figure 1A and 1B). To understand whether this result was influenced by the assay system used, a subset of validating and non-validating TCRs was introduced into primary human peripheral blood CD8^+^ T cells. Using CD137 expression as a measure of TCR signaling in human CD8^+^ T cells, a high level of agreement between the two assay systems was observed (Figures 1C and 1D). Comparison of validating and non-validating TCRs based on tcrdist3 sequence similarity suggests that sequence conservation may be used to predict reactivity for the majority of YLQ TCRs but less so for GLC-reactive TCRs (Figure 1E). Jointly, these analyses provide evidence that the data quality of VDJdb is poor, potentially limiting its value for TCR specificity model training and evaluation.

**Figure 1.**
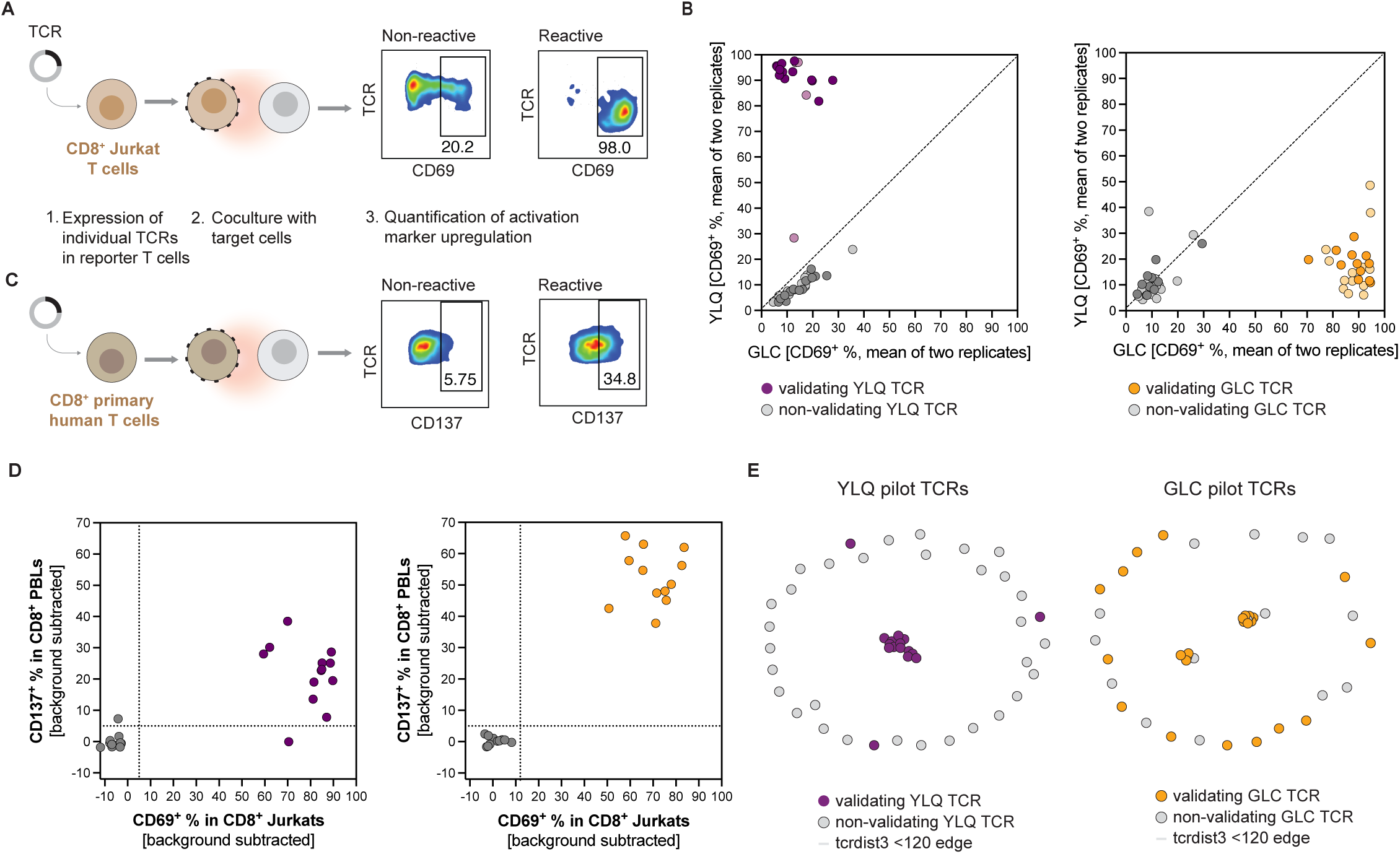
Exploration of data quality in VDJdb. (A) Assay setup and representative flow cytometry data depicting CD69 expression on TCR-transduced CD8αβ⁺ TCR-null Jurkat cells following coculture with either YLQ-or GLC-expressing HLA-A*02:01^+^ B cells. (B) Scatterplot depicting CD69 expression on TCR-transduced CD8⁺ TCR-null Jurkat cells upon exposure to either YLQ-or GLC-expressing HLA-A*02:01^+^ B cells. Dots represent individual TCRs. Dots with darker shading indicate TCRs that were also evaluated in primary CD8⁺ T cells, as shown in (C and D). (C). Assay setup and representative flow cytometry data depicting CD137 expression on TCR-transduced primary CD8⁺ T cells following coculture with either YLQ-or GLC-expressing HLA-A*02:01^+^ B cells. (D). Scatterplot comparing CD69 expression on TCR-transduced CD8⁺ TCR-null Jurkat cells and CD137 expression on TCR-transduced primary CD8⁺ T cells, following coculture with HLA-A*02:01^+^ B cells expressing the indicated epitopes. Dots represent individual TCRs. Background CD69/CD137 signal observed upon co-culture with HLA-A*02:01^+^ B cells expressing irrelevant epitopes was subtracted. (E) tcrdist3 distance-based network formed by YLQ– and GLC-annotated TCRs evaluated in (A-D). Grey nodes represent non-validating TCRs, colored nodes represent validating TCRs in (A,B). Nodes are connected by edges when TCRs represented by those nodes have a tcrdist3 distance below 120.

### Scalable synthetic TCR library assembly and characterization

To allow the evaluation of a major fraction of TCR-pMHC entries in VDJdb or in future datasets, we set out to create a high-throughput synthetic system for TCR generation and functional screening. Value of such a synthetic system is formed by its sole requirement for TCR sequence and epitope sequence information as input. In addition, use of such a synthetic system removes variation in T cell (dys)function as a confounder. Assembly of large synthetic TCRαβ libraries requires the precise combination of defined TCRα and β chains into gene segments of approximately 1.8 kilobase that are difficult to synthesize in a cost-efficient and hence scalable manner. Previous approaches have aimed to increase scalability by arrayed TCR assembly of TRAV– and TRBV-encoding vectors in combination with TCR-specific CDR3α/β-Jα/β-encoding oligonucleotides^1,2^, but throughput remains limited by the requirement for individual custom oligonucleotides. As an alternative, recent work has described a scalable TCR assembly approach that relies on oligonucleotide pools that encode all required CDR3α/β-Jα/βs segments in combination with re-usable TRAV– and TRBV-encoding vector stocks in a one-pot reaction^26^. However, relatively low sequence accuracy and imbalanced composition of resulting TCR libraries make it difficult to ensure sensitive screening of all TCRs within libraries, especially for those TCRs that are present at low frequencies.

Building on these approaches, we reasoned that a scalable robotics-based TCR assembly method could be developed that combines high assembly accuracy with the scalability of oligonucleotide pools (Figures 2A and S1A). In brief, to accurately demultiplex oligonucleotide pools to achieve assembly of unique TCRs in a TCR library of interest, we selected (see Methods) and assigned primers from a set of highly orthogonal (non-interacting) primers^27^ to wells in plates, allowing the generation of PCR-amplified CDR3α-Jα and CDR3β-Jβ segments of defined TCRs in each well. Joining of the resulting fragments by Golden Gate assembly with TRAV and TRBV fragments and constant regions could then be used to yield full-length TCRβ-P2A-TCRα gene segments encoded by viral vectors (for details see Methods). To ensure the intended ligation during Golden Gate assembly, highly orthogonal four-base overhangs using a comprehensive ligation fidelity dataset^28^ were utilized (Figure S1B and S1C).

**Figure 2.**
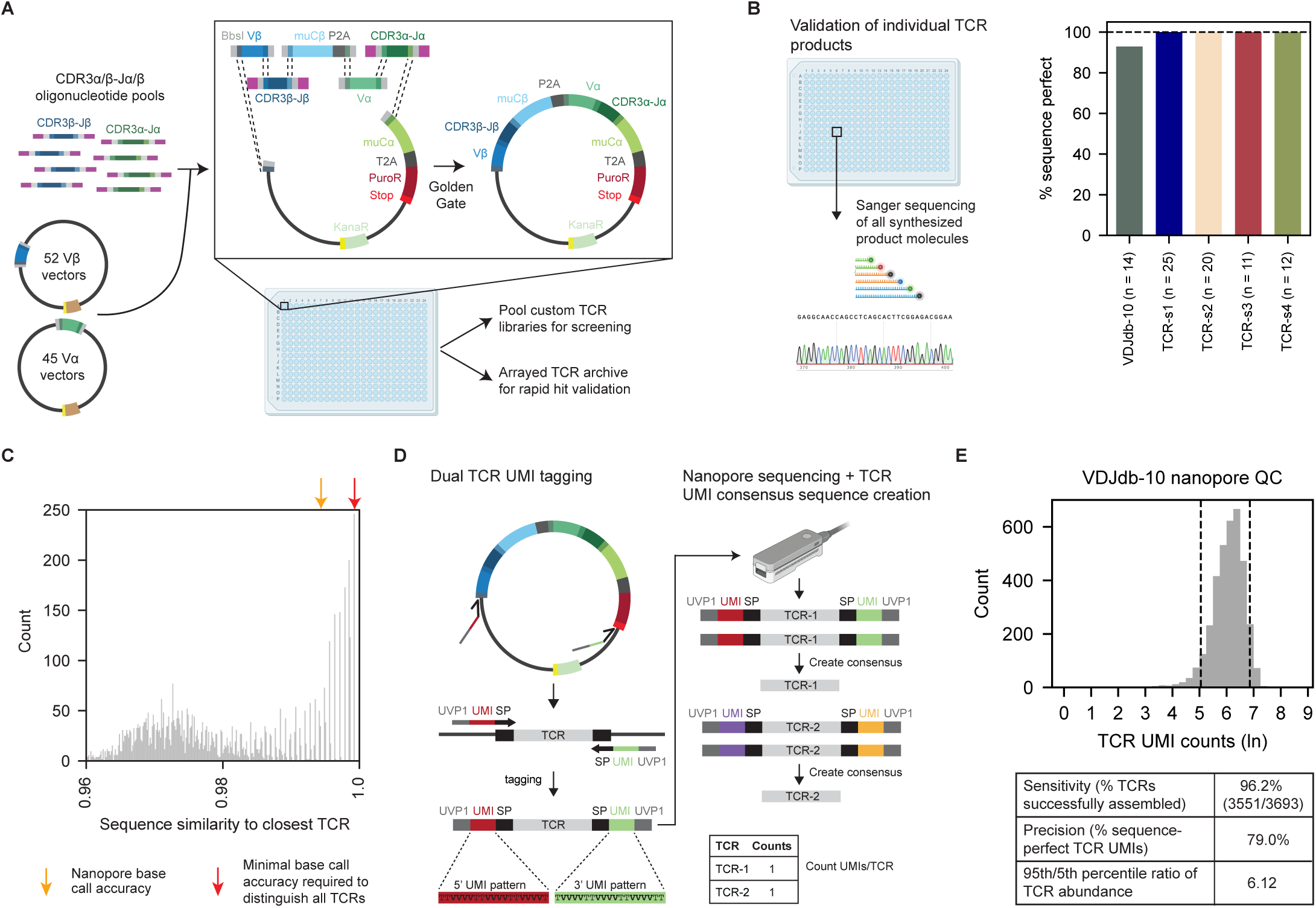
Scalable synthetic TCR library assembly and characterization. (A) Schematic overview of synthetic TCR library assembly platform (see Figure S1A for details). (B) Percentage of full-length sequence-perfect (i.e., exact matching sequences) synthetic TCRs, as evaluated using Sanger-sequencing of bulk TCR assembly products. Indicated numbers of assembly products of the VDJdb-10 (3,693 TCRs), TCR-s1 (3,510 TCRs), TCR-s2 (983 TCRs), TCR-s3 (1,000 TCRs), and TCR-s4 (2,892 TCRs) libraries were evaluated. (C) Sequence identity between each TCR in the VDJdb-10 library and its closest match, based on pairwise comparisons. Identities were calculated by self-aligning all TCRs using minimap2^47^, defined as BLAST identity: exact nucleotide matches at the same position, counting each consecutive gap separately. The orange arrow marks the average raw base-calling accuracy of Oxford Nanopore Technologies (ONT), while the red arrow indicates the minimum sequence accuracy threshold above which all TCRs in the VDJdb-10 library are distinguishable. (D) Schematic of ONT sequence unique molecular identifier (UMI)-based error correction method to sequence unique full-length TCR molecules at the accuracy required to distinguish all TCRs. (see Figure S2A-B for details of ONT-sequencing DNA library preparation and error correction bioinformatics). Unique synthetic TCR molecules are tagged with UMIs at both ends. ONT sequencing-reads originating from the same unique TCR molecule are identified by UMI binning and used to create an accurate consensus sequence by medaka polishing. Resulting consensus sequences are aligned to the reference sequence list to allow counting of unique full-length TCR molecules. (E) Histogram of TCR UMI counts of the VDJdb-10 library and corresponding count QC summary statistics.

To evaluate the performance of this high-throughput TCR assembly method we created a diverse set of 12,078 MHC class I– and II-restricted TCRs in five synthesis runs at a cost of €2.40/ TCR: i) all 3,693 TCRs in VDJdb that are annotated as reactive with one of the top 10 most densely sampled epitopes (“VDJdb-10” library), and ii-v) the TCR-s1 (3,510), TCR-s2 (983), TCR-s3 (1,000), and TCR-s4 (2,892) libraries that cover a range of tumor-derived and designed TCRs. To estimate the percentage of sequence-perfect TCR assemblies, defined as TCR sequences with correctly joined TCRβ-P2A-TCRα fragments with no mutations, insertions, or deletions, we performed full-length TCR Sanger sequencing of 82 randomly selected individual TCR assembly products. Sequence analysis of these bulk TCR assembly products from each of the five libraries showed that >98% of TCRs (81 out of 82 TCRs evaluated) was successfully assembled as sequence-perfect DNA molecules (Figure 2B). To extend this analysis of full-length TCR sequences to entire TCR library assemblies, we developed an Oxford Nanopore Technologies (ONT) full-length TCR sequencing method with sequencing error correction^29^ to be able to count sequence-perfect unique TCR molecules (Figure 2C, 2D, S2A, S2B). In brief, the read length of ONT sequencing data in principle enables accurate estimation of Vα–CDR3α-Jα and Vβ–CDR3-Jβ chain pairings. However, due to high sequence similarity between different TCR chains (Figure 2C), its raw base-call accuracy is insufficient to confidently call correctly paired or mispaired TCRs, nor detect mutations or indels, and comprehensive QC metrics of TCR libraries have to date not been available. To create a QC pipeline with single base error detection capacity, we designed an ONT sequencing method that utilizes unique molecular identifiers (UMIs) to bin reads, thereby enabling creation of highly accurate consensus sequences for individual TCR molecules (Figure 2D, S2A and S2B). Resulting consensus sequences allow for precise identification of sequence variations including mutations and indels, as well as correct and incorrect chain pairings, as shown by a ‘leave one plate out’ repeat assembly experiment (Figure S2C). Analysis of unique TCR molecules in each of the five TCR libraries showed successful assembly of 96.9% (range: 96.2% – 98.6%) of all TCRs, with on average 74.2% sequence-perfect unique TCR molecules (Figure 2E and S2D). Importantly, in line with the strategy for saturation amplification of individual CDR3α/β-Jα/β segments (see Methods), library composition was highly uniform, with abundance between the 5th and 95th percentiles in the five TCR libraries varying on average only 5.6-fold (Figure 2E and S2D). Taken together, these data describe a TCR assembly method that is highly accurate at the single-molecule and single-nucleotide level. In addition, the UMI-based ONT approach for full-length TCR sequencing also provides a strategy to distinguish reactive and non-reactive TCR sequences with single base resolution in genetic screens (see below).

### Pooled functional genetic screening of the VDJdb-10 TCR library

To generate a high-confidence dataset for TCR specificity modeling, we set out to validate the reactivity of the VDJdb-10 TCR library against the YLQ and GLC epitopes (Figure 3A), building on a recently developed functional genetic screening approach to identify tumor-reactive TCRs in human tumor-infiltrating lymphocyte (TIL) samples^26^. Following introduction of the VDJdb-10 TCR library in TCR-null CD8^+^ Jurkat T cells, library composition and uniformity was confirmed by ONT sequencing of TCR^+^ cells (Figure 3B). Subsequently, the VDJdb-10 Jurkat library was exposed to either YLQ-or GLC-expressing target cells, and reactive T cells were isolated for ONT sequencing by cell sorting of CD69^+^ cells (Figure S3A). Using TCR abundance in the CD69^+^ cell pools as a measure of TCR signaling, reactivity could be confidently called for 171 out of 421 (40.6%) YLQ-annotated TCRs and for 134 out of 193 (69.4%) GLC-annotated TCRs (Figure 3C). Importantly, lack of claimed TCR reactivity was not explained by TCR abundance in the VDJdb-10 Jurkat library (Figure S3B and S3C), nor by collapse of non-*01 TRAV and TRBV alleles to *01 alleles (Figure S3D). Furthermore, TCR reactivity as measured by pooled functional screening was highly consistent with the results obtained by the initial arrayed analyses, with a concordance of 96% (Figure S3E). Interestingly, evaluation of features that were associated with lack of claimed reactivity revealed a substantial variation in data quality between studies (Figure S3F). Furthermore, the fraction of samples obtained from donors with a recent or ongoing COVID-19 infection across studies was highly predictive of the fraction of truly reactive TCRs as identified here (Figure 3D), suggesting that antigen-driven expansion may help enrich for truly reactive TCRs. Importantly, VDJdb confidence scores showed a low specificity and also low sensitivity in identifying truly reactive TCRs for both epitopes (Figure 3E and 3F), highlighting the lack of reliable quality control metrics in currently used TCR-pMHC databases. Collectively, these functional screening data demonstrate that the key property of TCRs for predictive model development, i.e., their ability to signal upon encounter of endogenously presented antigen, is lacking in a large fraction of annotated VDJdb TCRs. In addition, we provide a functionally validated VDJdb database (“TCRvdb”) for model evaluation, and a standardized evaluation platform to rapidly expand this dataset to other TCR – antigen collections.

**Figure 3.**
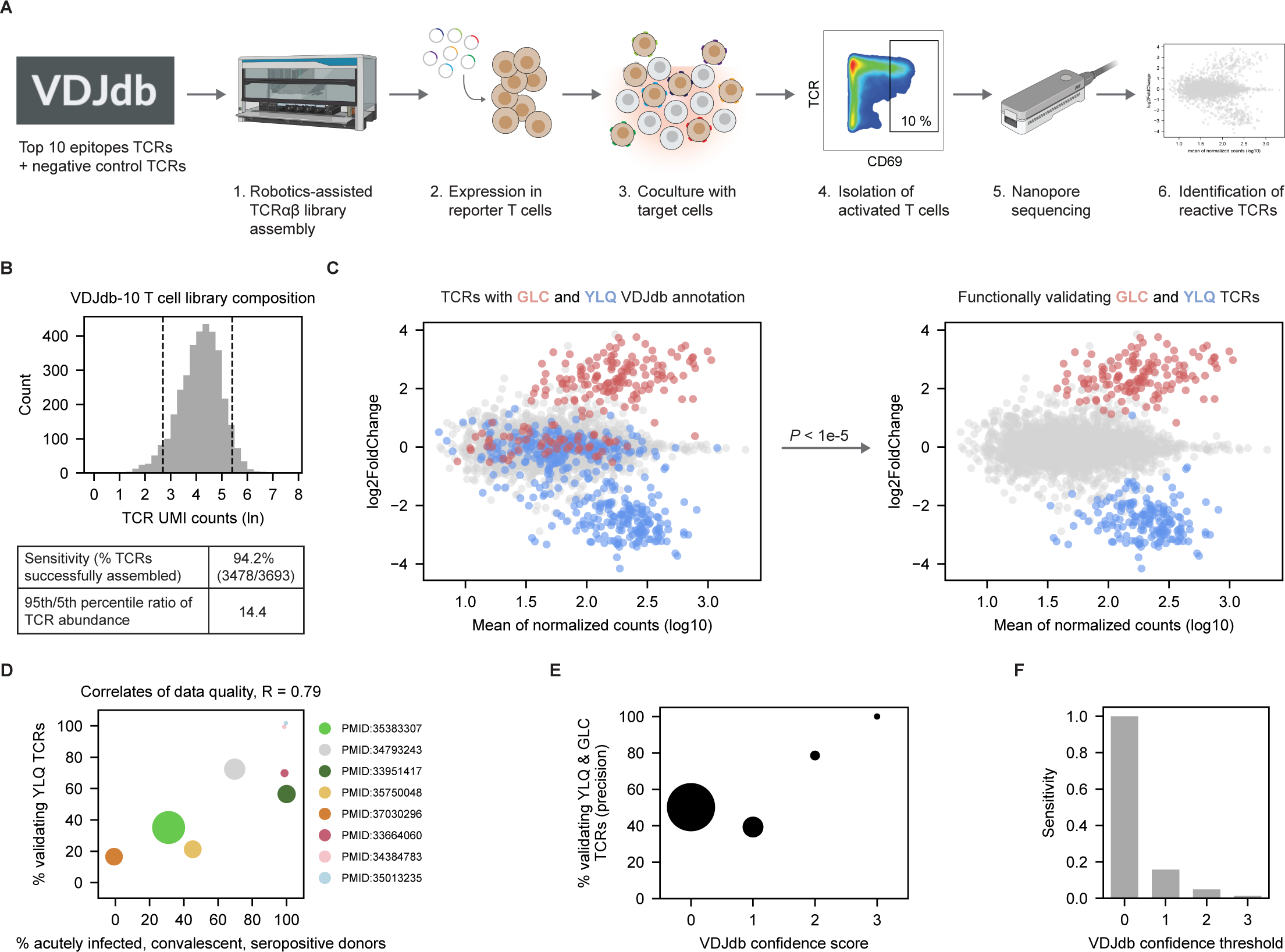
Pooled functional genetic screening of the VDJdb-10 TCR library. (A) Schematic overview of pooled functional genetic screening approach. The VDJdb-10 TCR library was introduced into CD8αβ^+^ TCR-null Jurkat cells. VDJdb-10-expressing Jurkat cells were exposed to HLA-A*02:01^+^ B cells expressing either the YLQ or GLC epitope and reactive T cells were isolated using CD69^+^ cell sorting. Abundance of individual VDJdb-10 TCRs in the CD69^+^ cell fractions was determined by full-length TCR UMI ONT-sequencing. Reactive TCRs were then identified by their relative abundance in the different CD69^+^ cell fractions. (B) Histogram of TCR UMI counts of the VDJdb-10 library in Jurkat T cells and corresponding count QC summary statistics. (C) MA plot of fold changes, representing the relative abundance of TCR UMI counts in the YLQ epitope exposed and GLC epitope-exposed CD69^+^ cell fractions over the mean normalized TCR UMI count across CD69^+^ cell fractions. Dots represent individual TCRs. Left MA plot: dots are colored by VDJdb annotation for reactivity against GLC (red) or YLQ (blue). Right MA plot: significantly enriched (*P* value < 1e-5) dots are colored. *P* values were calculated using the DESeq2^48,49^ Wald test and adjusted for multiple comparisons. (D) Scatterplot depicting the percentage of validating YLQ-reactive TCRs in individual studies as a function of the percentage of samples from acutely infected, convalescent, or SARS-CoV-2 seropositive donors. In all studies, TCRs were identified based on pMHC multimer binding. In those cases where clonotype-level donor information was available (the majority of studies) the fraction of TCR clonotypes originating from acutely infected, convalescent, and seropositive donors was determined. Dot size indicates the number of TCRs contributed by each study (range 1-191 TCRs). (E) Plot depicting the percentage of validating YLQ– and GLC-annotated TCRs (i.e., precision) at the indicated VDJdb confidence scores. Dot size indicates the total number of annotated TCRs at each confidence score threshold (range 4-512 TCRs). (F) Barplot depicting the percentage of all validating YLQ– and GLC-annotated TCRs that is retained (i.e., sensitivity) when using the indicated thresholds of VDJdb confidence scores as a minimum quality threshold.

### Functionally validated VDJdb improves TCR-pMHC reactivity model evaluation

To understand whether the use of TCRvdb alters our interpretation of the precision and sensitivity of existing TCR-pMHC reactivity prediction models, we evaluated the tcrdist3^30^, STAPLER^19^, and Alphafold3^14^ models (Figure 4A). tcrdist3 exploits sequence similarity to group TCRs with similar sequences at defined thresholds. While, by its nature, tcrdist3 cannot be trained to predict epitope reactivity, sequence similarity to one or multiple user-defined ‘anchor’ TCRs with known epitope reactivities can be utilized to identify additional TCRs that may recognize the same epitope. In contrast to tcrdist3, STAPLER is a supervised language model that was trained to predict the epitope reactivity of TCRs, using the (frequently incorrect, see Figure 1 and 3) reactivity labels provided by VDJdb (see Methods). Finally, Alphafold3 was utilized as a general protein complex structure prediction model that has not explicitly been trained to predict the epitope reactivity of TCRs. To predict TCR reactivity from the AlphaFold3 output, we developed the TCRbridge model, which uses AlphaBridge^31^ to combine AlphaFold3’s three per-residue predicted confidence metrics at the TCRα–peptide and TCRβ–peptide interfaces.

**Figure 4.**
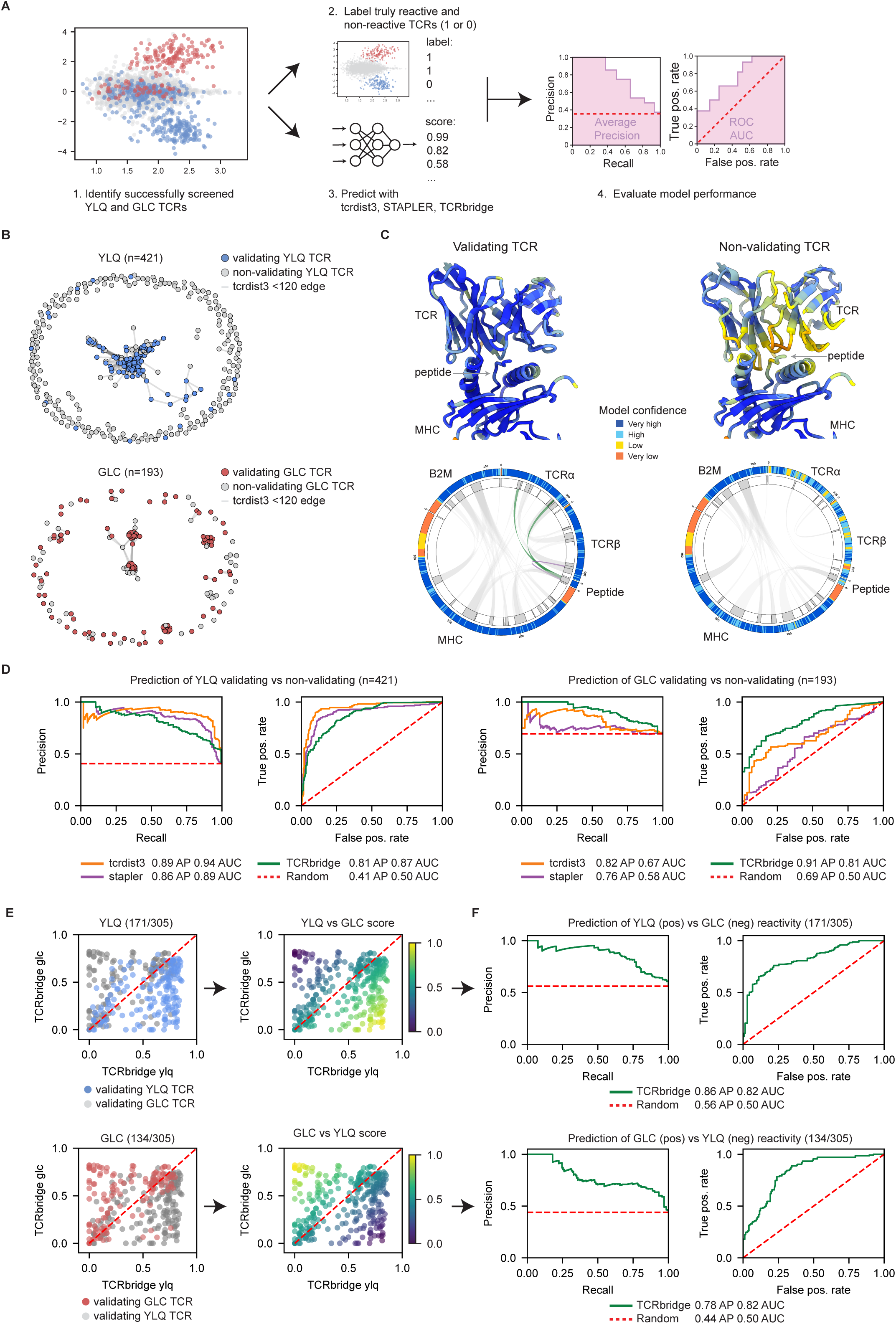
TCRvdb improves TCR-pMHC reactivity model evaluation. (A) Schematic overview of prediction model evaluation pipeline using validated VDJdb (TCRvdb). Validating and non-validating TCRs are identified using the functional genetic screening results as depicted in Figure 3C. tcrdist3, STAPLER, and TCRbridge are then evaluated on their ability to distinguish identified validating and non-validating TCRs. (B) tcrdist3 distance-based network formed by YLQ– and GLC-annotated TCRs. Grey nodes represent non-validating TCRs, colored nodes represent validating TCRs. Nodes are connected by edges when TCRs represented by those nodes have a tcrdist3 distance below 120. (C) AlphaFold3 (AF3) predicted TCR-pMHC structure complexes of a validating (top-left) and non-validating (top-right) YLQ-annotated TCR. Structures are colored by AF3’s confidence in the placement of each atom within its local environment (pLDDT; predicted Local Distance Difference Test). The corresponding AlphaBridge^31^ circos plots of the validating (bottom-left) and non-validating (bottom-right) predicted structure complexes show pLDDT for TCRα, TCRβ, peptide, MHC, and B2M (outer ring), along with confident interfaces identified by AlphaBridge (inner ring and ribbons). Ribbons representing confident interfaces between TCRα and peptide are colored green, between TCRβ and peptide are colored purple, and all other interfaces are shown in grey. Note the presence of confident interactions between TCR and peptide and TCR and MHC in the left model but not the right model. (D) Comparison of tcrdist3 (orange), STAPLER (purple), and TCRbridge (green) performance in predicting whether YLQ-annotated (left) and GLC-annotated (right) TCRs do or do not validate in functional genetic screening. Note that the labels identified in Fig. 3 were not included in the development of either model. Left: Precision-Recall curve showing the fraction of validating pairs of all pairs (i.e. precision) at the indicated recall (sensitivity) levels. Right: Receiver Operating Characteristic (ROC) curve showing ability of the indicated models to rank validating pairs above non-validating pairs, as measured as the true positive rate (sensitivity) versus the false positive rate (1 − specificity) across prediction thresholds. Average Precision (AP; area under the Precision-Recall curve) summarizes overall model performance on the Precision-Recall curve, with values closer to 1.0 indicating better precision. Area under the ROC curve (AUC) summarizes overall model performance in ranking validating pairs relative to non-validating pairs across prediction thresholds, with values closer to 1.0 indicating a higher ability to distinguish validating from non-validating pairs. Dashed red lines show random performance. (E) Left: Scatterplots comparing TCRbridge-predicted confidence scores of validating YLQ and GLC TCRs for structure predictions in combination with either their cognate or non-cognate epitopes. Top-left plot depicts scores for YLQ TCRs in blue. Bottom-left plot depicts scores for GLC TCRs in red. Top-right and bottom-right plots depict data for the same TCRs but colored by relative confidence score for both predictions. Top-right: higher scores indicate preference for YLQ over GLC. Bottom-right higher scores indicate preference for GLC over YLQ. Relative scores were computed by calculating the scalar projections of each TCR prediction onto a vector contrasting the two epitopes (either (1, –1) for YLQ or (–1, 1) for GLC), followed by min-max normalization to scale values from 0 (favoring the non-target epitope) to 1 (perfectly aligned with the target epitope). (F) Performance of TCRbridge in distinguishing cognate (positive) from swapped (negative) validating TCR-epitope pairs using the predicted relative score from (E). Precision-recall and ROC curves show model performance for validating YLQ (top) and GLC (bottom) TCRs. Precision-recall curves show the fraction of cognate epitope pairs of total pairs (precision) at the indicated levels of recall (sensitivity). ROC curves reflect the model’s ability to rank cognate epitope pairs relative to swapped pairs. As explained in (D), AP and AUC summarize overall model performance. Dashed red lines represent random performance.

Importantly, analysis of the output of the functional genetic screening data (Figure 3) using these three models showed that each of these displayed a moderate to substantial capacity to distinguish validating and non-validating TCRs, but with important nuances (Figure 4 and S4). First, while tcrdist3 showed a high capacity to distinguish truly YLQ-reactive TCRs from non-validating TCRs with claimed YLQ-reactivity, performance in distinguishing validating and non-validating TCRs with claimed GLC reactivities was modest (Figure 4B and S4A). Like tcrdist3, the masked language-based sequence model STAPLER showed some performance in distinguishing truly reactive from non-reactive TCRs, particularly for the YLQ epitope (Figure S4A). Notably, this performance was observed in spite of the fact that STAPLER was fine-tuned using a dataset in which both the validating and non-validating TCRs as identified here were all labeled as positives. We conclude that noise in datasets can only efficiently be identified with these sequence-based models in cases in which validating and non-validating TCRs are sufficiently distinct in sequence space. In contrast, TCRbridge demonstrated substantial performance in distinguishing reactive from non-reactive TCRs for both the YLQ and GLC epitopes (Figures 4C, 4D, and S4A), and outperformed tcrdist3 and STAPLER in identifying GLC-reactive TCRs (Figure 4D). Predictive capacity was also obtained when using a single confidence measure (Figure S4B) or interface confidence between TCR subunits and MHC (Figure S4C and S4D). These data suggest that, despite not being trained to predict TCR-pMHC reactivity in any way, AlphaFold3 has a superior capacity to correctly predict pMHC reactivity in case of more complex TCR repertoires.

In future prediction tasks, TCRs that share the same HLA restriction element (e.g., HLA-A*02:01) may be compared for their potential reactivity with a given peptide-MHC. However, the HLA restriction element of the non-validating TCRs as described here is unknown and may not be HLA-A*02:01. Furthermore, because of the bias towards HLA-A2*02:01-restricted TCRs in the AlphaFold3 training dataset, TCRbridge could potentially assign higher confidence to complexes that contain HLA*02:01-restricted TCRs. To understand whether the performance of TCRbridge on the TCRvdb dataset could be due to a learned HLA-A*02:01-restriction class imbalance, we predicted interfaces between all validated GLC-reactive TCRs and either the cognate GLC-HLA-A*02:01 complex or the control YLQ-HLA-A*02:01 complex (for which these TCRs are experimentally confirmed true negatives, see Figure 3). Likewise, interfaces were predicted between all validated YLQ-reactive TCRs and either the cognate YLQ-HLA-A*02:01 complex or the control GLC-HLA-A*02:01 complex. Notably, TCRbridge still assigned meaningful confidence scores in this setting, indicating that model performance was not due to a learned HLA-A*02:01 restriction class imbalance (Figure 4E, 4F, S4E, and S4F). In contrast, this ability to distinguish TCR-pMHC complexes that differ by only the 9-mer epitope sequence was not seen with non-validating TCRs for either epitope (Figure S4G), further highlighting how TCRbridge performance estimates are dependent on data quality.

## Discussion

When low quality datasets are used to evaluate predictive models, this may lead to an underestimate of model performance, especially when these models have not seen, and hence been biased by, these data during training. We here demonstrate that data quality is a significant concern for the VDJdb database that has been the starting point of TCR-pMHC prediction evaluation efforts^21^. We also share a technology to synthetically recreate TCR collections and validate their function in a standardized fashion. Finally, we exploit this platform to yield validated datasets for two epitopes and then use these datasets to demonstrate that performance of established sequence– and structure-based prediction pipelines is higher than assumed^19,32,33^.

The VDJdb database was created through the joint effort of many research groups that examine T cell-mediated immune responses against pathogens, cancer antigens and other antigen classes. In line with its bottom-up nature, contributors have significant freedom with respect to assay systems (e.g., using different types of MHC multimers, using fresh or restimulated cell populations, using material from acutely infected or healthy donors) and thresholds used to call reactive TCR-pMHC pairs. Equally important, unlike in the field of structural biology, where well-established metrics^34,35^ are used to unambiguously describe data quality, it has been difficult to define standardized validation metrics that comprehensively describe data quality of TCR-pMHC pairs. We set out to evaluate previously proposed TCR-pMHC pairs in a well-defined experimental setting to create harmonized and validated datasets for model development. A key component in this effort has been the use of immortalized synthetic systems for both antigen and TCR expression, in order to remove the variability associated with finite biological samples. Importantly, as the goal of TCR-pMHC prediction efforts is to predict productive T cell signaling upon encounter of antigen-expressing cells, we define TCR reactivity based on T cell activation marker expression, rather than pMHC binding. Output of these pooled functional genetic screens was well-correlated with arrayed pilot screening of the same TCRs using distinct functional readouts. In addition to activation marker expression as used here, productive signaling through the T cell receptor may also be analyzed by e.g., antigen-induced proliferation or degranulation, or using distinct cell systems. We note though that the threshold for productive TCR signaling can depend on e.g., costimulatory signals, and it will thus be important to calibrate such alternative systems.

For both the datasets evaluated here, the fraction of non-validating TCRs was remarkably high. Notably, the fraction of TCR-pMHC pairs that could be experimentally validated varied substantially between studies, and, in case of SARS-CoV-2, was positively correlated with the fraction of samples from donors with recent or acute infections. We conclude that MHC multimer-based isolation of T cells from healthy donors frequently results in the isolation of TCRs that fail to signal upon antigen encounter, and this may either reflect random assay noise or may point towards the occurrence of TCRs that bind the relevant pMHC but do not productively signal, an issue that remains to be addressed.

For all three prediction models evaluated we identified an – in some cases substantial – capacity to distinguish those TCR-pMHC pairs that do or do not pass experimental validation. However, only for AlphaFold3 substantial performance was observed in the more demanding situation in which valid and non-valid TCRs show high sequence similarity (Fig. 4). Interestingly, recent evaluations of AlphaFold2 PAE confidence-based approaches observed an only modest performance in TCR-pMHC prediction for well-studied epitopes in VDJdb^32,33^. Based on the data presented here, we speculate that the seemingly modest performance of these models may in fact reflect their capacity to also partly distinguish validating and non-validating TCR-pMHC complexes. The PDB version that was used to train AlphaFold3 included structures of two TCR-YLQ-HLA-A2 complexes (7N1F and 7N6E) and one TCR-GLC-HLA-A2 complex (3O4L) (training cutoff date: 30 September 2021^14^). Prediction confidence of TCRbridge for these three structures did not exceed average prediction confidence (Figure S4F), arguing against data leakage as a driver of the observed performance. Notably, TCRbridge not only performed well in distinguishing validating and non-validating TCRs, but also in assigning epitope reactivity when analyzing two sets of TCRs that share the same HLA-A*02:01 restriction element. The TCRbridge reactivity score that we describe here combines the three AlphaFold3 confidence measures at the TCR-pMHC interface and appears to yield slightly better performance as compared to the use of single confidence measures (Figure S4B). With the upcoming availability of validated TCR reactivity datasets for a larger number of epitopes, it will be attractive to utilize supervised machine learning models trained on the full range of features extractable from AlphaFold3 to identify those feature combinations that are most predictive of TCR-pMHC reactivity across epitopes. By the same token, it will also be attractive to use these data to investigate alternative biophysical scoring approaches, for instance extending TCRbridge with force field interaction energies^36^, as force fields were recently shown to outperform supervised models trained on AlphaFold2-derived peptide, MHC, and TCR structural features in predicting the recognition of unseen epitopes^33^. Next to these approaches to optimize extraction of the relevant information from structure predictions that are obtained using the generic version of structure-based algorithms, it will be valuable to use large validated TCR reactivity datasets to directly fine-tune AlphaFold3 model parameters for TCR specificity prediction, as was recently demonstrated for the prediction of peptide-MHC binding^37^.

The scalable ‘sequence-synthesis-screen’ approach presented here may be utilized for a number of purposes. First, in line with the strategy chosen here, the approach may be used to assess the function of TCR-pMHC pairs that have been described by researchers in different fields over the past decades. Second, the synthetic systems described here should form a valuable quality control step for ongoing large-scale data collections efforts that e.g., aim to describe neoantigen-specific T cell reactivity across cancer patient cohorts, or that aim to describe the TCR repertoire for model antigens to saturation. Finally, the synthetic nature of the technology makes it well-suited to explore reactivity beyond the TCRs that are observed in natural samples. For instance, the predictive capacity of TCR-pMHC specificity models may be evaluated by the design of TCRs with pre-specified criteria, and experimental testing of thousands of predicted TCRs may then be utilized to understand features of predictive success and failure. Output from all three efforts should contribute to the validated dataset initiated here. To support its use in model development, the TCRvdb dataset will be hosted on https://github.com/schumacherlab/TCRvdb as a resource to the community.

## Resource Availability

### Lead Contact

Requests for further information and resources and reagents should be directed to W. Scheper and T. Schumacher.

### Materials Availability

The VDJdb-10 TCR plasmid library is made available for academic research upon request.

### Data and Code Availability

All ONT sequencing data will be deposited in the National Center for Biotechnology Information’s Sequence Read Archive. The TCRtoolbox python package is available at: https://github.com/schumacherlab/TCRtoolbox. The ONT-TCRconsensus package is available at: https://github.com/schumacherlab/ONT-TCRconsensus. The validated VDJdb database is available at: https://github.com/schumacherlab/TCRvdb. The TCRbridge python package is available at: https://github.com/schumacherlab/TCRbridge. Code use to generate figures in this paper is available at: https://github.com/schumacherlab/schumacherlab_publications_code.

## Acknowledgments

This work was supported by the Dutch Cancer Society Young Investigator Grant (no. 2020-1/12977) (to W.S.); ZonMw Translational Research Program 2 (no. 446002001) (to W.S.) and by the Louis Jeantet prize and Stevin Award (to T.N.S.). This work was also created as part of the MATCHMAKERS team, supported by the Cancer Grand Challenges partnership financed by CRUK (CGCATF-2023/100003) and the National Cancer Institute (OT2CA297206). RLB was supported by Oncode Accelerator (grant number NGFOP2201) and by the ScreeninC infrastructure (KWF 12539). Research at the Netherlands Cancer Institute is supported by institutional grants from the Dutch Cancer Society and the Dutch Ministry of Health, Welfare and Sport. We thank the Research High Performance Computing facility of the Netherlands Cancer Institute for providing and maintaining computation resources. We thank the flow cytometry core at the Netherlands Cancer Institute for technical support. We would like Benoit Nicolet and other members of the Schumacher group for scientific input.

## Author contributions

T.N.S, W.S., and M.Me. conceived and designed the study. M.Me., W.S., J.U., B.P.Y.K., T.N.S., J.G., B.M., Ž.M., S.d.P., M.S., and R.L.B. developed the TCR assembly method. Ž.M., S.d.P., R.V., A.G., M.Me., Y.W., M.Mu., designed and performed T cell activation and functional genetic screening experiments. M.Me., J.G., Ž.M., J.U., and B.P.Y.K. developed the full-length TCR ONT-sequencing method. B.P.Y.K., M.Me., D.Á.S., Ž.M., M.I., A.P., J.T., E.M., performed data analysis and modelling. M.Me., T.N.S. and W.S., wrote the manuscript. T.N.S. and W.S. supervised the project. All authors reviewed and revised the manuscript.

## STAR Methods

### Experimental model and subject details

#### Cell lines

FLY-RD18 packaging cells (Sigma) were used for production of retroviral supernatants. TCRα^-^/β^-^ CD8αβ^+^ Jurkat cells were described previously^38^. Primary CD8^+^ T cells were isolated from peripheral blood mononuclear cells (PBMCs) of healthy donors (Sanquin, the Netherlands). Immortalized B cell lines were generated by retroviral introduction of Bcl-6 and Bcl-xL into B cells isolated from donor PBMCs, as previously described^26^.

### Method details

#### Antibodies

The following antibodies were used for flow cytometry: anti-CD4-FITC (clone RPA-T4, BD Biosciences, dilution 1:200); anti-CD19-FITC (clone 4G7, BD Biosciences, dilution 1:200); anti-CD8α-APC (clone SK1, BioLegend, dilution 1:50); anti-CD69-BV421 (clone FN50, BioLegend, dilution 1:50); anti-CD137-BV421 (clone 4B4, BioLegend, dilution 1:50). PE-conjugated anti-mouse TCRβ constant domain (clone H57-597, BD Biosciences, dilution 1:150). Live/Dead Fixable Near-IR Dead Cell Stain (Thermo Fisher Scientific), or DAPI was used to identify live cells. Data from flow cytometry experiments were acquired using FACSDiva software (version 8.0.2, BD Biosciences) and analyzed using FlowJo (version 10.7.1, BD Biosciences).

#### Robotics-assisted Golden Gate cloning of synthetic TCR libraries

Individual synthetic TCRs were assembled and cloned into the pMX retroviral expression plasmid in separate wells of 384-well plates. Briefly, Golden Gate cloning was used to join the CDR3α-Jα, CDR3β-Jβ, TRAV, TRBV, and murine TRBC-P2A (muTRBC-P2A) fragments into a pMX retroviral vector that also encodes murine TRAC and the puromycin N-acetyltransferase resistance gene (Figure S1A). To enable PCR amplification of desired CDR3α/β-Jα/β fragments from complex oligonucleotide pools in individual wells of 384-well plates, 384 unique and highly orthogonal forward and reverse primer combinations (‘well-specific’ primers) were used (Supplementary Table 2). Separate unique primer combinations (‘plate-specific’ primers) were utilized to enable PCR amplification of 384-well-plate-specific sub-pools of CDR3α/β-Jα/β fragments from larger oligonucleotide pools (Supplementary Table 3). Individual CDR3α/β-Jα/β fragments were amplified by two rounds of targeted PCR: (i) amplification of 384-well-plate-specific oligonucleotide sub-pools using plate-specific primers, followed by (ii) amplification of individual CDR3α-Jα and CDR3β-Jβ fragments using the assigned 384-well orthogonal primer combinations. As a means to reduce the differential abundance of individual CDR3α/β-Jα/β fragments in synthetic oligonucleotide pools (previously observed to contribute to non-uniformity of final TCR libraries^26^), PCR reactions in the second round were carried out into plateau phase. The TRAV and TRBV genes required for assembly of each unique TCR were then added to individual wells from TRAV and TRBV plasmid stocks (Supplementary Table 4). Finally, a Golden Gate reagent mix that also contains muTRBC-P2A and pMX-muTRAC-T2A plasmids was added to the wells, and all fragments were joined by Golden Gate assembly. All process steps, further detailed below, were carried out using largely automated dispensers to ensure scalability of synthetic TCR assembly.

##### Well-specific CDR3α-Jα and CDR3β-Jβ fragment generation

In order to enable PCR amplification of desired CDR3α/β-Jα/β fragments from complex oligonucleotide pools in individual wells of 384-well plates with high specificity, we tested the PCR efficiencies of a collection of highly orthogonal primers^27^ and assigned the best performing primers to 384 unique forward and reverse primer combinations (well-specific primers). Additional efficient primer pairs were chosen to enable PCR amplification of plate-specific sub-pools of oligonucleotides from larger pools. For each 384-well plate, a sub-pool of plate-specific CDR3α/β-Jα/β fragments was PCR-amplified by combining 0.230 ng CDR3α/β-Jα/β oligonucleotide pool (Twist Bioscience), 0.2 mM dNTPs (Sigma; 11969064001), 0.02 U/µL Q5 polymerase (NEB; M0493L), 0.25 µM of both forward and reverse plate-specific primers, and 1x Q5 polymerase buffer in a total volume of 140 µL, followed by incubation in a thermocycler (30 s at 98 °C, 16 cycles of 10 s at 98 °C, 20 s at 65 °C and 35 s at 72 °C, followed by 2 min at 72 °C). Amplified CDR3α/β-Jα/β fragment sub-pools were purified using the NEB Monarch DNA Column Kit (T1130L) and PCR product yield was measured using an Agilent Bioanalyzer 2100 (DNA 7500 kit, 5067-1506).

Using a Hamilton/I.DOT dispenser, well-specific CDR3α/β-Jα/β fragments were amplified in 384-well plates from amplified CDR3α/β-Jα/β fragment sub-pools by combining 0.15 ng PCR-amplified sub-pool with 0.2 mM dNTPs, 0.02 U/µL Q5 polymerase and 0.75 µM forward and reverse primers, and 1x Q5 polymerase buffer in a total volume of 2.67 µL in each well. Forward and reverse primer combinations were pre-added to 384-well-plates using a Hamilton Microlab STAR 384 liquid handler system, and PCR amplification mix was dispensed into the 384-well-plates using an I.DOT dispenser (Dispendix, BICO), followed by incubation in a thermocycler (30 s at 98 °C, 39 cycles of 10 s at 98 °C, 20 s at 63.5 °C and 25 s at 72 °C, followed by 2 min at 72 °C). Amplified well-specific CDR3α/β-Jα/β fragments were diluted 5.3-fold with nuclease-free water (Invitrogen; #4387936) using a Hamilton/I.DOT dispenser and PCR yield was routinely verified for a small selection of wells using an Agilent Bioanalyzer 2100 (DNA 7500 kit). Lastly, PCR products were diluted another 13.4-fold with nuclease-free water (to approximately 220 pg/µL) using a Hamilton/I.DOT dispenser in order to reduce the concentrations of PCR reagents that inhibit downstream Golden Gate reactions.

##### Golden Gate assembly of synthetic TCRs

To prevent mis-ligation of TCR fragments during Golden Gate assembly, we first designed a highly orthogonal set of four-base overhangs using a comprehensive overhang ligation fidelity^28^ dataset (Figure S1B and S1C). Golden Gate assembly of synthetic TCRs was performed by automated dispensing of 8.55 ng pMX-muTRAC-T2A retroviral plasmid, 14 ng of muTRBC-P2A plasmid, 11.88 ng of both TRAV and TRBV plasmids (Twist Bioscience), 840 pg of PCR-amplified well-specific CDR3α/β-Jα/β fragments, 20 U/µL T4 ligase (NEB; M0202M), 0.5x BbsI (Thermo Fisher Scientific; #FD1014), and 1x T4 ligase buffer in each well of a 384-well plate, in a total volume of 4.75 µL, followed by incubation of 384-well plates in thermocyclers (50 cycles of 10 min at 37 °C and 10 min at 16 °C, followed by 10 min at 55 °C and 10 min heat inactivation at 65 °C).

Following Golden Gate assembly, reaction products were pooled using only part [2 µL] of the product in each well, to thereby retain a repository of individual TCRs. To purify DNA from pooled Golden Gate products, DNA was precipitated with isopropanol (1:1 volume ratio), GlycoBlue Coprecipitant (100 ng/µL), and 0.05 M NaCl. Precipitated DNA was washed twice with 80% ethanol and eluted in 25 µL nuclease free water. Purified Golden Gate product pools were transformed at >1,000x coverage into electrocompetent Endura *E.coli* (60242-2, Biosearch Technologies) by electroporation (10 uF, 600 ν, 1800 V), using a BioRad Gene Pulser Xcell. Bacteria were subsequently grown overnight at 37 °C in LB medium containing kanamycin. The use of distinct antibiotic resistance genes in the used plasmids (Fig. S1A) enables counter-selection of remaining muTRBC-P2A, TRAV and TRBV plasmid material (Fig. S1A). 384-well plates containing individual TCR archives were stored at –20 °C for use in, for instance, downstream characterization of selected TCRs.

#### Preparation of long read TCR sequencing libraries

##### Dual-UMI-tagging and nested PCR amplification

For each individual TCR in a library, we tagged 50,000 (plasmid DNA) or 100 (gDNA) template molecule copies (Figure S2A). Each template molecule was tagged with terminal UMIs by PCR, using primers nanop_GSP_umi_Fw and nanop_GSP_umi_Rev (Supplementary Table 5). Both primers contain a universal primer site for downstream amplification and a patterned UMI consisting of 18 `V` nucleotides (all nucleotides except `T`) and 14 patterned ‘anchor’ `T` nucleotides, yielding a combined complexity of 1.5 × 10¹⁷ UMI sequences.

To estimate the average rate of UMI collisions per template molecule (i.e., the average fraction of distinct molecules that are indistinguishable due to being tagged with the same UMIs), we used an alternative formulation of the birthday problem^39^. For this calculation, the effective number of UMIs was estimated using the 3-error Hamming bound, as 3-errors conservatively corresponds to clustering UMIs with VSEARCH^40^ at 93% identity. Based on this model, the estimated maximal average UMI collision rate per template molecule is 2.3 × 10^-10^ for plasmid (19 individual TCRs in the largest VDJdb-10 TCR similarity bin × 50,000 copies) and 4.6 × 10^-13^ gDNA (19 individual TCRs × 100 copies) templates.

Dual-UMI-tagging PCR reactions were performed by combining the calculated amount of template DNA (50,000 plasmid copies corresponding to 0.178 pg or 100 gDNA copies corresponding to 700 pg × the number of unique TCRs in the library) with 0.5 µM forward and reverse primers, 1x NEBNext HiFi 2x PCR mastermix (NEB; M0541L) in a total volume of 15 µL (plasmid) or 100 µL per 67 ng input gDNA, followed by incubation in a thermocycler (3 min at 98 °C, 2 cycles of 30 s, 90 s at 72°C and 90 s at 72°C, followed by 5 min at 72°C). After dual-UMI tagging, tagging primers were removed using 0.6x AMPure XP beads (Beckman Coulter), and reaction products were washed twice with 80% ethanol, and eluted in 20 µL (plasmid), or 20 µL per 200 µL required dual-UMI-tagging reaction volume (gDNA), nuclease-free water.

Dual-UMI-tagged full-length TCR molecules were further amplified by two rounds of nested PCR using the primers in Supplementary Table 5. In the first PCR, complete dual-UMI-tagged reaction products were PCR-amplified using 0.5 µM nanop_UVP_umi_1_Fw and nanop_UVP_umi_1_Rev primers and 1x NEBNext HiFi 2x PCR mastermix in a total volume of 96 µL. The reaction mixture was incubated in a thermocycler (1 min at 98 °C, 5 cycles of 15 s at 98 °C, 25 s at 67 °C and 90 s at 72 °C, immediately followed by 5 cycles of 15 s at 98 °C, 25 s at 72 °C and 90 s at 72 °C, and a final 2 min at 72 °C). PCR products were purified using 0.6x AMPure XP beads, washed twice with 80% ethanol, eluted in 20 µL nuclease-free water. Amplification products were then further PCR-amplified by combining complete round one reaction products with 0.5 µM nanop_UVP_2_Fw and nanop_UVP_2_Rev primers and 1x NEBNext HiFi 2x PCR mastermix in a total volume of 96 µL, followed by incubation in a thermocycler (1 min at 98 °C, followed by a set number of cycles of 15 s at 98 °C, 25 s at 70 °C and 90 s at 72 °C, and 2 min at 72 °C). Here, the number of PCR cycles was based on the amount of initial plasmid or gDNA template that was required to tag all TCR template molecules with UMIs at sufficient coverage. PCR products were purified using 0.6x AMPure XP beads, washed twice with 80% ethanol, eluted in 13 µL nuclease-free water, and product concentrations were measured using an Agilent Bioanalyzer 2100 (DNA 7500 kit).

##### ONT long read TCR sequencing

Full-length TCR amplicons were prepared for ONT sequencing using the Native Barcoding Kit 24 V14 protocol (Oxford Nanopore Technologies, SQK-NBD114.24). In brief, 200 fmol of amplicon DNA was end-prepped and dA-tailed using NEBNext Ultra II End Repair/dA-Tailing Module (NEB; E7546), followed by ligation of barcodes from the Native Barcoding Expansion kit using Blunt/TA Ligase (NEB; M0367). Barcoded samples were pooled and subjected to native adapter ligation with the NEBNext Quick Ligation Module (NEB; E6056). Resulting libraries were quantified using a Qubit dsDNA HS Assay (Thermo Fisher) and sequenced at 2,000-2,500x coverage, using a R10.4.1 flow cell on MinION or PromethION sequencing devices.

#### Expression of individual TCRs and TCR libraries in reporter T cells

For expression in Jurkat T cells, TCRs were retrovirally transduced into TCRα^−^/TCRβ^−^ CD8αβ^+^ Jurkat T cells. In brief, retroviral supernatant was produced by transfection of FLY-RD18 packaging cells with pMX plasmid DNA encoding indicated TCR libraries or individual TCRs using Lipofectamine 3000 Reagent (Thermo Fisher Scientific). After 48 h, supernatants were collected, filtered and centrifuged (2,000 g for 90 min) in 24-well plates precoated with RetroNectin (Takara). CD8^+^ TCRα^−^/TCRβ^−^ Jurkat cells were infected by incubation on virus-coated plates for 21 hours. For the production of TCR libraries, Jurkat cells were transduced at infection rates of less than 5% to ensure single retroviral integrations in the majority of transduced cells. After 72 h, transduction efficiency was assessed by cell staining with anti-mouse TCRβ constant domain antibody and flow cytometry. TCR-expressing Jurkat cells were selected by culture in the presence of 2.5 µg/ml puromycin and then 5.0 µg/ml puromycin for 48 h each. Subsequently, TCR-transduced cells were isolated by staining with anti-mouse TCRβ constant domain antibody and cell sorting on a FACSAria Fusion (BD Biosciences) to achieve at least 80% purity. Sorted Jurkat cells were cultured in RPMI 1640 medium supplemented with 10% FBS (Sigma-Aldrich) until use in TCR discovery screens. TCR library coverage of at least 250x was maintained during all steps.

For TCR expression in primary T cells, CD8^+^ T cells were isolated from healthy donor PBMCs using the CD8^+^ T Cell Isolation Kit (Miltenyi Biotec) and stimulated with anti-CD3/anti-CD28 Dynabeads (Life Technologies) in RPMI 1640 medium supplemented with 10% human AB serum (Life Technologies), penicillin–streptomycin (Roche) and 150 U mL^−1^ IL-2 (Proleukin, Novartis) for 48 h. 24-well plates precoated with RetroNectin (Takara) were coated with retrovirus by centrifugation (2,000 g for 90 min) of retroviral supernatants, followed by incubation of stimulated CD8^+^ PBMCs on these retrovirus-coated plates for 21 h. Transduction efficiency was measured after 72 h by cell staining with anti-mouse TCRβ constant domain antibody and flow cytometry. Transduced cells were selected by culture in the presence of 2.5 µg/ml puromycin for 48 h, with subsequent addition of new IL-2 and medium every 3–4 days.

#### Generation of antigen-expressing B cell lines

The HLA-A*02:01-restricted `GLCTLVAML` (GLC) EBV-derived and `YLQPRTFLL` (YLQ) SARS-Cov2-derived epitopes were introduced into an HLA-A*02:01^+^ Bcl-6/Bcl-xL-immortalized B cell line by lentiviral transduction. In brief, lentiviral supernatant was produced by transfection of 293T LentiX HEK cells (Takara) with a lentiviral plasmid encoding mKelly1 and either the GLC or YLQ epitope, and psPAX2 and pGALV^41^ helper plasmids, using Lipofectamine™ 3000 Reagent (Thermo Fisher Scientific). After 48 h, supernatants were collected, filtered and centrifuged (2,000 g for 90 min) in 24-well plates precoated with RetroNectin (Takara). Immortalized B cells were infected by incubation on virus-coated plates for 21h. Transduction efficiency was measured after 72 h by analysis of mKelly1 expression by flow cytometry. Transduced B cells were subsequently selected with 2.5 – 5.0 µg/mL puromycin for 48 h and cultured in the presence of irradiated (55 Gy) CD40L^+^ mouse L cells in IMDM medium (Gibco) supplemented with 10% heat-inactivated FBS (Sigma-Aldrich), penicillin-streptomycin and 50ng/mL IL-21 (BioLegend)) until further use. B cells received new irradiated CD40L^+^ L cells every 10 days, and new medium and IL-21 every 3-4 days.

#### T cell activation assays

Reactivity of T cells transduced with single TCRs was analyzed by flow cytometry after overnight coculture with target cells in U-bottom 96-well plates at an effector-to-target ratio of 1:1. T cells incubated in the absence of target cells with or without phorbol 12-myristate 13-acetate (50 ng/mL; Sigma-Aldrich) and ionomycin (1 µg/mL; Sigma-Aldrich), served as positive and negative controls, respectively. Cells were stained with IR-Dye and anti-mouse TCRβ constant domain antibody, in combination with anti-CD8 and anti-CD137 (when using primary T cells) or with anti-CD69 (when using Jurkat cells) antibodies, and subsequently analyzed by flow cytometry. Coculture with irrelevant epitope-expressing B cells was used to determine background T cell activation levels.

#### Functional TCR reactivity screens

TCR discovery screens were performed as described previously for TIL-derived TCRs^26^. In brief, functional genetic screens were performed by incubating VDJdb-10 TCR library-expressing CD8^+^ Jurkat T cells with B cells expressing the indicated epitopes at an effector-to-target ratio of 1:1. Subsequently, Jurkat cells with the highest 10% of CD69 expression were isolated using a FACSAria Fusion cell sorter and TCR abundance was determined by nanopore sequencing, as described above. Functional genetic screens were performed in triplicate, with a TCR library coverage of 500x in all experiments. Reactive TCRs and non-reactive TCRs were identified from nanopore sequencing data as described below.

### Data analysis and Statistical Analysis

#### Creation of the VDJdb-10 dataset

The VDJdb database was downloaded from the antigengenomics/vdjdb-db GitHub page on the 19^th^ of May 2023 and was built directly from submissions using provided instructions. The vdjdb_full.txt file was filtered to only contain human CD8^+^ T cell-derived TCRs and MHC-I binding epitopes. In addition, the database was filtered to only contain unique and complete TCR-epitope annotations (filtering on TRAV, TRBV, TRAJ and TRBJ genes, CDR3α and CDR3β amino acid sequences, and epitope amino acid sequence). All entries derived from 10X Genomics data from the Application Note – A New Way of Exploring Immunity (v1 Chemistry) were removed (https://www.10xgenomics.com/resources/datasets), as prior work provided evidence for the low quality of this dataset^42^. This resulted in 6,450 paired TCR-pMHC entries. Fifty duplicate TCRs were removed because the annotated epitopes were at >2 edit distance of each other. One TCR entry was removed because of a missing HLA-gene annotation. 28 epitope entries were removed because their HLA-gene annotation did not match the HLA annotated in an experimentally verified dataset of peptide HLA allele binders^43^. 64 entries were removed because they contained an HLA annotation that does not occur in the IMGT HLA allele database^44^ (i.e., “A*08:01”, “A*02:256”, and “B*51:193”). The final parsed VDJdb dataset, in which TCR-pMHC pairs that would be most likely to contain errors have been removed, contained 6,307 entries.

To create the VDJdb-10 TCR library from the final parsed VDJdb dataset, we extracted all 3,251 TCRs annotated as reactive to the top 10 most densely sampled epitopes in VDJdb: `GILGFVFTL`, `YLQPRTFLL`, `TTDPSFLGRY`, `NLVPMVATV`, `SPRWYFYYL`, `CINGVCWTV`, `LLWNGPMAV`, `GLCTLVAML`, `LTDEMIAQY`, and `ATDALMTGF`. Additionally, we included 549 negative control TCRs annotated as reactive to epitopes that have only one or two TCR entries in VDJdb, collectively spanning 466 unique epitopes.

#### Design of TCR library assemblies

We developed a Python package, called `TCRtoolbox`, to design and prepare TCR library assemblies, requiring only the TRAV, TRAJ, TRBV and TRBJ gene names and CDR3α and CDR3β amino acid sequences as input. As a first step, the package translates TRAV, TRAJ, TRBV, and TRBJ gene names into standardized IMGT format. `TCRtoolbox` subsequently trims the TRAJ and TRBJ gene amino acid sequences from their N-terminus to the first amino acid of the J gene end motif (either `FGXG`, `WGXG`, `XGXG`, or `FXXG`)⁴⁵. The trimmed J genes are then joined to their corresponding CDR3α/β regions to generate CDR3α/β-Jα/β amino acid sequences. Resulting sequences are codon optimized using a custom codon optimizer within `TCRtoolbox` to eliminate type IIS BbsI recognition sites, four-base overhangs (Figure S1C), Kozak sequences, primer binding sites, repeats longer than four nucleotides, and terminator motifs (`TTTTT` or `AAAAA`). Codon optimization enforces a global GC content between 35–60%, with a relaxed range of 25– 70% allowed only for sequences that cannot otherwise be optimized. Sequences failing these criteria are filtered out. All VDJdb-10 TCR CDR3α/β-Jα/β sequences were successfully codon optimized.

`TCRtoolbox` subsequently assigns individual TCRs to wells of 384-well plates, together with unique primer pairs that were selected from a set of highly orthogonal primers. For each TCR that is assigned to a defined well of a 384-well plate, the corresponding primer nucleotide sequences are added to the 5’ and 3’ ends of the codon-optimized CDR3α/β-Jα/β nucleotide sequences by `TCRtoolbox`. Resulting nucleotide sequences are written to an Excel worksheet that may directly be used to obtain oligonucleotide pools from a commercial supplier. In parallel, `TCRtoolbox` assigns TRAV and TRBV genes from each TCR to each well. These genes are stored as reusable plasmid stocks in 2D-barcoded tubes, currently covering the *01 alleles of all TRAV and TRBV genes (Supplementary Table 4). For seamless joining with CDR3 sequences, TRAV and TRBV amino acid sequences are trimmed from their C-termini up to the cysteine residue at position 104 (Cys104, IMGT numbering). `TCRtoolbox` automatically keeps track of TRAV and TRBV stock volumes. `TCRtoolbox` also writes a Hamilton automatic pipetting instruction Excel sheet to generate an I.DOT dispenser source plate from the 2D-barcoded tubes, and I.DOT dispense instruction CSV file is written to add the required TRAV and TRBV genes from the resulting I.DOT source plate to the corresponding 384-wells for TCR assembly.

To check the constructed CDR3-J fragment sequences for design errors and TRAV and TRBV 384-well assignment before start of physical TCR assembly, we simulate cloning of each TCR *in silico* using `pydna`⁴⁶, modeling both restriction digestion, ligation, and PCR to yield the fully assembled TCR-encoding retroviral plasmid sequences. The resulting TCR amino acid sequence is then compared to the independently reconstructed amino acid sequence of the same TCR using custom TCR reconstruction software implemented in `TCRtoolbox`, as previously described¹⁹.

#### TCR UMI consensus creation and counting pipeline

ONT sequencing reads were base-called with dorado using the sup@v5.0.0 model with a `–-min-qscore` of 10: https://github.com/nanoporetech/dorado. To create high-accuracy full-length TCR UMI consensus reads that can be counted, we developed a Python package called `ONT-TCRconsensus`, with steps from^29,45^. All exact parameter settings and program versions used by this package are provided on its GitHub page. In this package, primers are first trimmed from raw base-called ONT reads using dorado, and trimmed reads with an expected error rate ^46^ (EE rate) >7%, as predicted from base-call qualities are removed using `VSEARCH`^40^. As it is not possible to unambiguously assign filtered raw base-called reads to reference TCRs that have highly similar sequences, we developed a pipeline that first assigns raw base-called reads to TCR similarity bins that contain reference TCRs with highly related sequences. This allows the subsequent creation of UMI-based consensus sequences with sufficient accuracy to unambiguously assign the consensus reads to specific TCR reference sequences. In this approach, the TCR library reference FASTA file is self-aligned using minimap2^47^ and TCRs with >93% sequence identity (as raw base-called reads maximally have a 7% EE rate) are clustered into ‘TCR similarity bins’.

Next, raw base-called reads are aligned to the TCR library reference with `minimap2`. Alignments are assigned to their respective TCR similarity bins based on the reference TCR that each raw base-called read aligns to. Within each TCR similarity bin, high-quality UMIs are detected on both ends based on their designed sequence pattern (see above) and correct UMI lengths. Subreads originating from the same unique TCR molecule are then identified by clustering the detected UMIs into UMI bins with `VSEARCH`. PCR chimeras, which create low-frequency novel UMI combinations by recombination, are removed by filtering out UMI bins with a subread coverage <4. Of note, `VSEARCH` UMI clustering within TCR identity clusters minimizes computational resource requirements. To validate TCR similarity binning, we confirmed that UMIs did not overlap between TCR similarity bins. The consensus sequence for each UMI bin is then generated from binned subreads using `medaka` polishing: https://github.com/nanoporetech/medaka. The resulting high-accuracy consensus sequences are aligned to the TCR reference library using `minimap2`. Alignments with a BLAST sequence identity (exact nucleotide matches at the same position, counting each consecutive gap separately) below the minimum required to distinguish all closest-identity TCR pairs in the reference are filtered, ensuring that only consensus sequences accurate enough to distinguish all TCRs in the library are counted. To validate the entire `ONT-TCRconsensus` pipeline, we confirmed that UMIs do not overlap between alignments to unique reference TCRs.

To calculate precision as % sequence-perfect reads, we filtered alignments below 1.0 BLAST identity (i.e., exact matching sequences). Finally, TCR UMI counts were generated by counting the number of unique UMIs of which the consensus sequences aligned to each reference TCR.

#### Reactive TCR identification using DESeq2

To identify reactive TCRs in pooled TCR screens, per-sample count tables were normalized and compared with `PyDESeq2`^48,49^ (version 0.5.0). Within the set of 3,428 VDJdb-10 TCRs (92.8% of all VDJdb-10 TCRs) that was detected across all YLQ and GLC triplicates, reactive and non-reactive TCRs were defined using the DESeq2 Wald test with an adjusted P < 1e-5. All default parameters were used, except `refit_cooks`, which was set to False.

#### tcrdist3 network analysis

Pairwise tcrdist3 distances were computed between all query TCRs, resulting in an n × n distance matrix, where n is the number of TCRs compared. Edges between TCR nodes were defined for any TCR pair with a TCRdist3 distance less than 120. This threshold was consistently used to generate all TCR networks. Force-directed layouts of the resulting networks were visualized using the `spring_layout` function from `NetworkX` (v2.7), with the layout parameter `k=0.15`.

#### Visualization of Alphafold3 predicted TCR-pMHC complex structures

AlphaFold3-predicted TCR–pMHC structures were visualized using ChimeraX, with residues colored according to AlphaFold3’s pLDDT confidence scores. Circus plots were generated using AlphaBridge. Protein interfaces, as defined by AlphaBridge, were colored green for TCRα-peptide interfaces, purple for TCRβ-peptide interfaces and grey for all other interfaces. The outer ring reflects the per residue model confidence (pLDDT) of the respective sequences.

#### TCR-pMHC reactivity modeling

##### tcrdist3

tcrdist3 distances were computed between each query TCR and all YLQ-or GLC-annotated TCRs, resulting in a 1 × (n−1) vector of distances. The nearest neighbors for each query TCR were identified by selecting the (n−1)/10 smallest distances (rounded to the nearest integer). A weighted average of these nearest neighbor distances was then computed, where smaller distances received higher weights. The final prediction score was the negative of this weighted average.

##### STAPLER

STAPLER was trained on the parsed VDJdb dataset of 6,307 TCR-pMHC entries. Training was performed using the training procedure outlined in^19^. Query TCRs were then predicted using the best model checkpoint from each fold, and the mean of the predicted scores across folds was used as the final prediction score for each query TCR.

##### pLDDT

AlphaFold3’s pLDDT confidence scores for predicted TCR-pMHC complex structures were parsed from the CIF files outputted by the AlphaFold3 inference implementation. Final pLDDT scores were computed as the mean pLDDT of the CDR3α, CDR3β and epitope that was extracted using the PDB parser of BioPython^50^ by taking b-factor score (pLDDT) of the Cα atom of each amino acid.

##### TCRbridge

TCR-pMHC complex structures were predicted using an implementation of the inference pipeline of AlphaFold3 (version 3.0.0): https://github.com/google-deepmind/alphafold3. AlphaBridge interface scores (ABi) were calculated for TCRα-peptide and TCRβ-peptide interfaces at an interface threshold of 0.4. If no interface was identified at this interface threshold, the TCRbridge prediction score was set to 0. The final TCRbridge prediction score was computed as the mean of the two TCRα-peptide and TCRβ-peptide ABi scores. Next to ABi scores, AlphaBridge also provides PDE, PMC and iptm_i scores for each interface.

#### Model performance evaluation using PRC and ROC-AUC

The scikit-learn^51^ implementation of `precision_recall_curve` was used to calculate the precision and recall for every threshold using prediction scores and labels. The `average_precision_score` was used to calculate average precision (AP), defined as the area under the precision-recall curve. Random average precision is defined as the number of positive samples divided by the total number of samples. The scikit-learn `roc_curve` implementation was used to calculate true positive and false positive rates for every threshold using prediction scores and labels. The `roc_auc_score` was used to calculate the area under the receiver operating characteristic curve (AUC). The random line is defined as 0.5 for all AUC curves. Labels can either be validating or non-validating TCRs (Figure 4D), or reactive or non-reactive TCR-pMHC pairs (Figure 4E, 4F, and S4G).

**Supplementary Figure 1.**
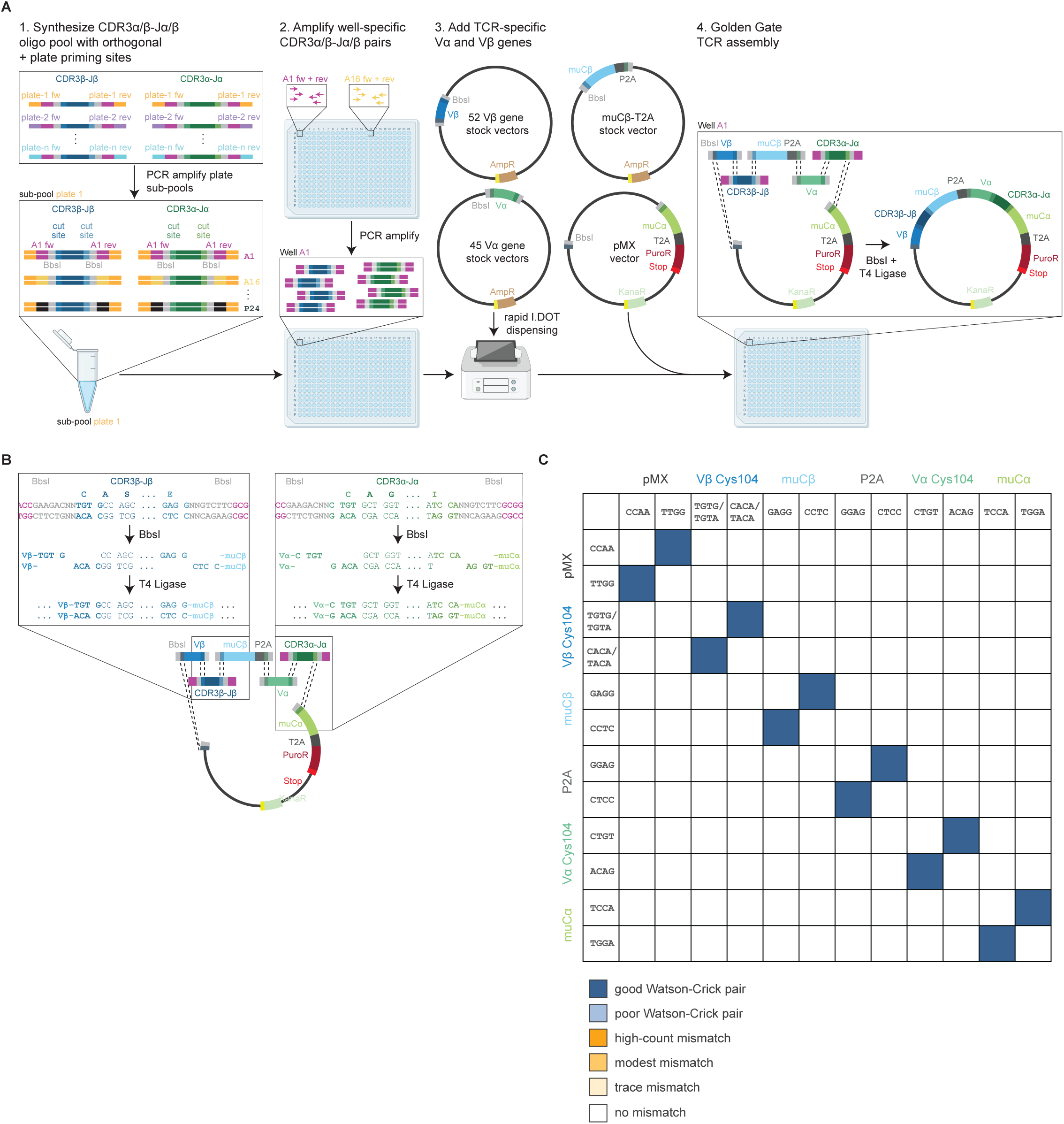
Scalable synthetic TCR library assembly platform. (A) Schematic overview of robotics-assisted arrayed synthetic TCR library assembly. All TCRs in a library are assembled in 384-well plate(s), with one unique TCR per well. CDR3α/β-Jα/β oligonucleotide pools are demultiplexed using highly orthogonal (non-interacting) primers, indicated in the oligonucleotide sequence in purple^27^. Each well is assigned a unique pair of orthogonal primers, enabling PCR amplification of DNA encoding the CDR3α-Jα and CDR3β-β segments of a unique TCR. Importantly, by amplifying all CDR3α-Jα and CDR3β-β segments individually until the PCR plateau phase, we can overcome the non-uniformity of the supplier-manufactured oligonucleotide pools, thereby generating highly uniform libraries. Robotics-assisted dispensing is then used to add the corresponding TRAV and TRBV fragments to each well. Golden Gate assembly is used to join the CDR3α-Jα and CDR3β-β, TRAV, and TRBV fragments with the murine TRBC-P2A (muTRBC-P2A), yielding a full-length TCRβ-P2A-TCRα gene segment encoded in the pMX retroviral vector. (B) Schematic overview of Golden Gate-based assembly of TRBV, CDR3β-Jβ, muTRBC, and TRAV, CDR3α-Jα, and muTRAC fragments. BbsI is used to generate user-defined four-base sticky overhangs that can be end-joined by T4 DNA Ligase. The four-base overhang that joins TRBV and CDR3β-Jβ is formed by the conserved Cys104 `TGT` codon at the start of the CDR3β-Jβ, together with a conserved `G` in all TRBV genes except TRBV20-1 and TRBV29-1, which instead contain a conserved `A` (conservation identified using sequence analysis of ∼89,000 TCRs^52^). The four-base overhang that joins TRAV and CDR3α-Jα is formed by the conserved Cys104 `TGT` codon at the start of the CDR3α-Jα, together with a conserved `C` nucleotide (conservation identified using the same sequence analysis of ∼89,000 TCRs). CDR3β-Jβ is joined to muTRBC using the `GAGG` overhang, and CDR3α-Jα is joined to muTRAC using the `TCCA` overhang. All other used four-base overhangs are shown in (C). (C) NEBridge Ligase Fidelity Viewer output for all four-base overhangs used to join fragments together forming a synthetic TCR. NEBridge Ligase Fidelity Viewer is based on a comprehensive T4 DNA ligase end-joining fidelity dataset^28^. Empty squares indicate no predicted mismatches.

**Supplementary Figure 2.**
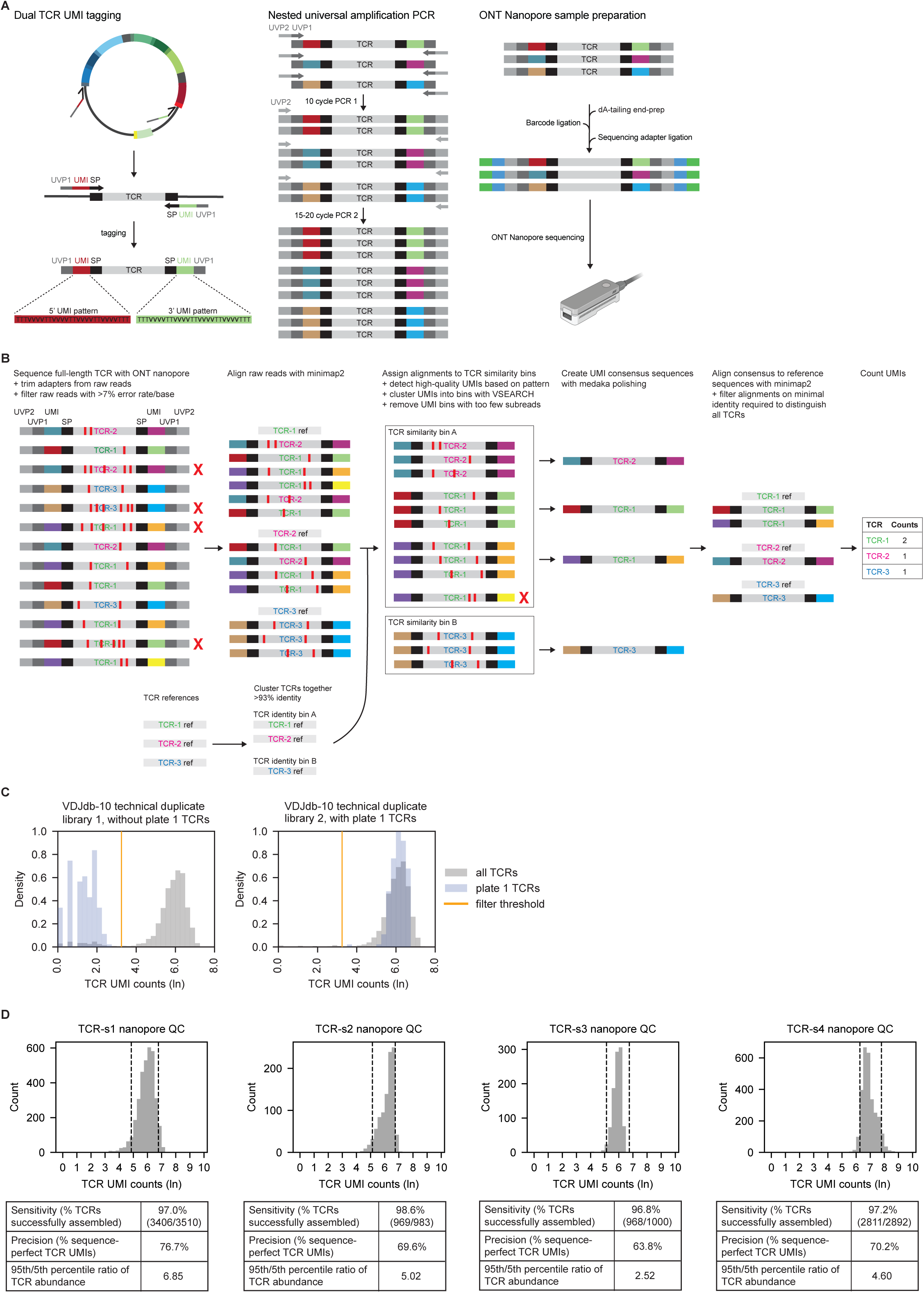
Nanopore-sequencing TCR library QC analysis. (A) Schematic overview of Oxford Nanopore Technologies (ONT) full-length TCR DNA sequencing library preparation. Unique TCR molecules are tagged on both ends with unique molecular identifiers (UMIs) using a 2-cycle PCR. UMI-tagged TCRs are then amplified by two nested PCRs. ONT barcodes and native sequence adapters are ligated, and resulting libraries are sequenced. (B) Schematic overview of the full-length TCR UMI sequence error correction pipeline. Due to the inherent base-calling errors of raw nanopore reads, TCRs with high sequence identity are difficult to distinguish. To address this, reference TCR sequences from the library are first self-aligned using minimap2, and sequences sharing greater than 93% identity are clustered into TCR similarity bins. Next, raw TCR nanopore reads are filtered using VSEARCH^40^ to exclude reads with more than a 7% expected error rate^46^ (note that this falls within the 93% identity threshold used for TCR reference sequence binning), and non-filtered raw reads are aligned to the reference TCR library using minimap2^47^. Each read is then assigned to a TCR similarity bin based on the reference sequence it aligns to. Within each TCR similarity bin, high-quality UMIs are identified and clustered using VSEARCH to group subreads derived from the same unique molecule into UMI bins. Importantly, UMI clustering is performed after alignment and binning by TCR identity, thereby reducing the chance of UMI collisions and computational load, as fewer UMI comparisons are required. Use of paired UMIs on both ends of the full-length TCR facilitates the detection and removal of PCR chimeras by filtering out UMI bins with insufficient subread support. A consensus sequence for each UMI bin is then created using the medaka polisher. Finally, high-accuracy consensus sequences are aligned to the reference TCR library using minimap2, and UMIs are counted for each unique TCR reference. (C) Histogram of TCR UMI count density for the VDJdb-10 plasmid library, when either synthesized with one plate omitted (left, plate 1 omitted) or synthesized in full (right). TCR UMI counts were normalized such that the area under each histogram integrates to 1, thereby representing the probability density of TCR UMI counts. The histogram for plate 1 TCRs is shown in purple, and all VDJdb-10 TCRs (including plate 1) are shown in grey. The orange vertical line marks the minimum TCR UMI count threshold used to filter out low UMI count noise. (D) Histogram of TCR UMI counts of the TCR-s1, TCR-s2, TCR-s3, and TCR-s4 libraries and their corresponding count QC summary statistics.

**Supplementary Figure 3.**
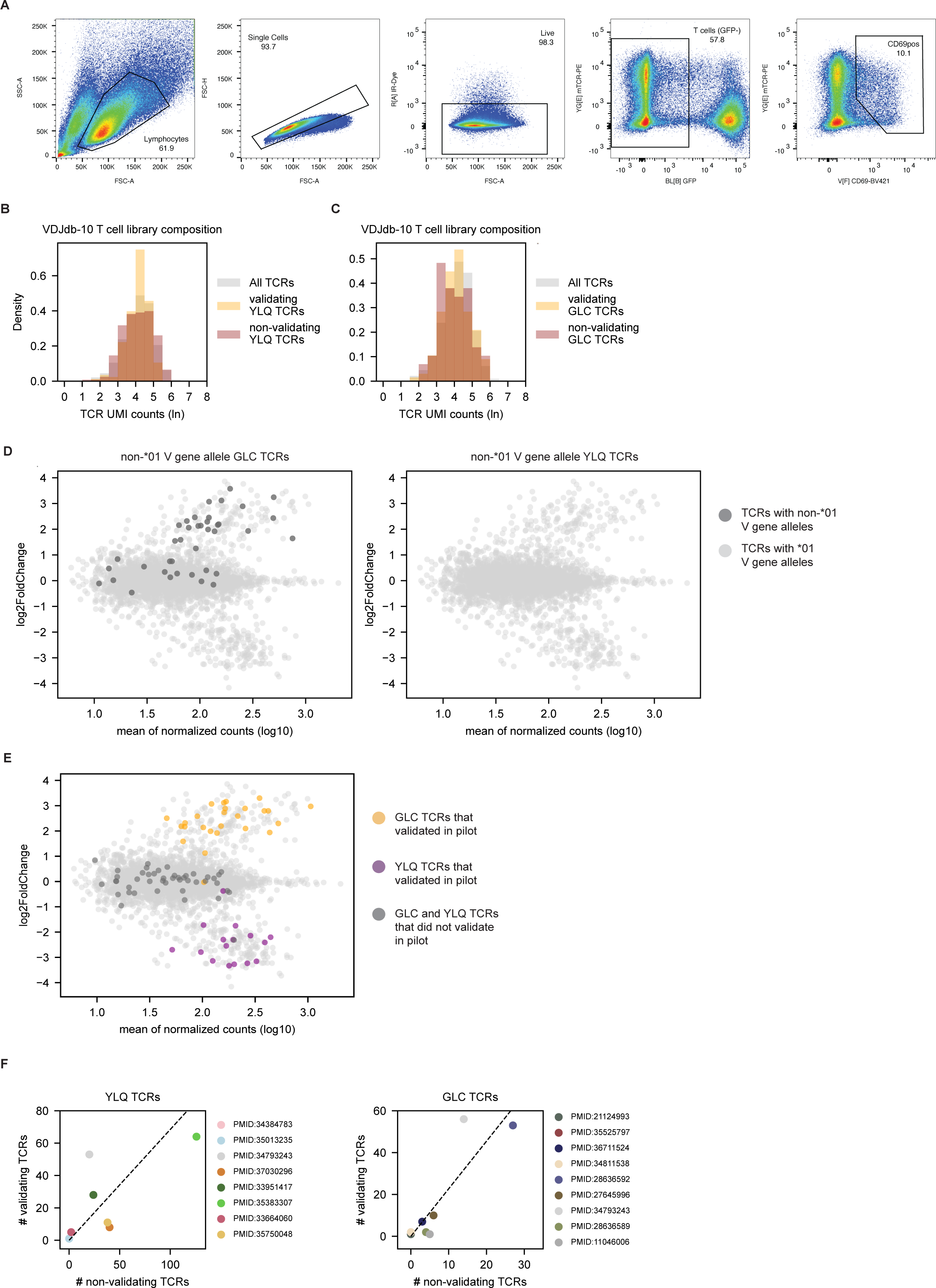
Pooled functional genetic screening of the VDJdb-10 TCR library. (A) Gating strategy for the isolation of CD69^+^ Jurkat T cells in the VDJdb-10 TCR library screen (relating to Figure 3A and 3C). (B) Histogram of TCR UMI count density of all VDJdb-10 TCRs (grey), and of validating (red) and non-validating (purple) YLQ TCRs in the screen of Figure 3C. TCR UMI counts were normalized such that the area under each histogram integrates to 1. (C) Histogram of TCR UMI count density of all VDJdb-10 TCRs (grey), and of validating (red) and non-validating (purple) GLC TCRs in the screen of Figure 3C. TCR UMI counts were normalized such that the area under each histogram integrates to 1. (D) MA plot from Figure 3C, with colors representing TCRs originally described with non-*01 (dark grey) or *01 (light grey) TRAV or TRBV alleles. Non-*01-alleles are collapsed to *01 in our synthetic TCR assembly method. Note that 10X Genomics single-cell sequencing workflow does not annotate non-*01 alleles, explaining the lack of non-*01 TRAV or TRBV alleles for all YLQ-annotated TCRs. (E) MA plot from Figure 3C, with dots colored by validation outcome in Figure 1A. Colors depict validating YLQ-annotated (purple) and GLC-annotated TCRs (yellow), and non-validating YLQ-annotated and GLC-annotated TCRs (both dark grey). (F) Scatterplots depicting the number of non-validating TCRs and validating TCRs contributed by individual studies. Studies are shown by PubMed unique study identification number.

**Supplementary Figure 4.**
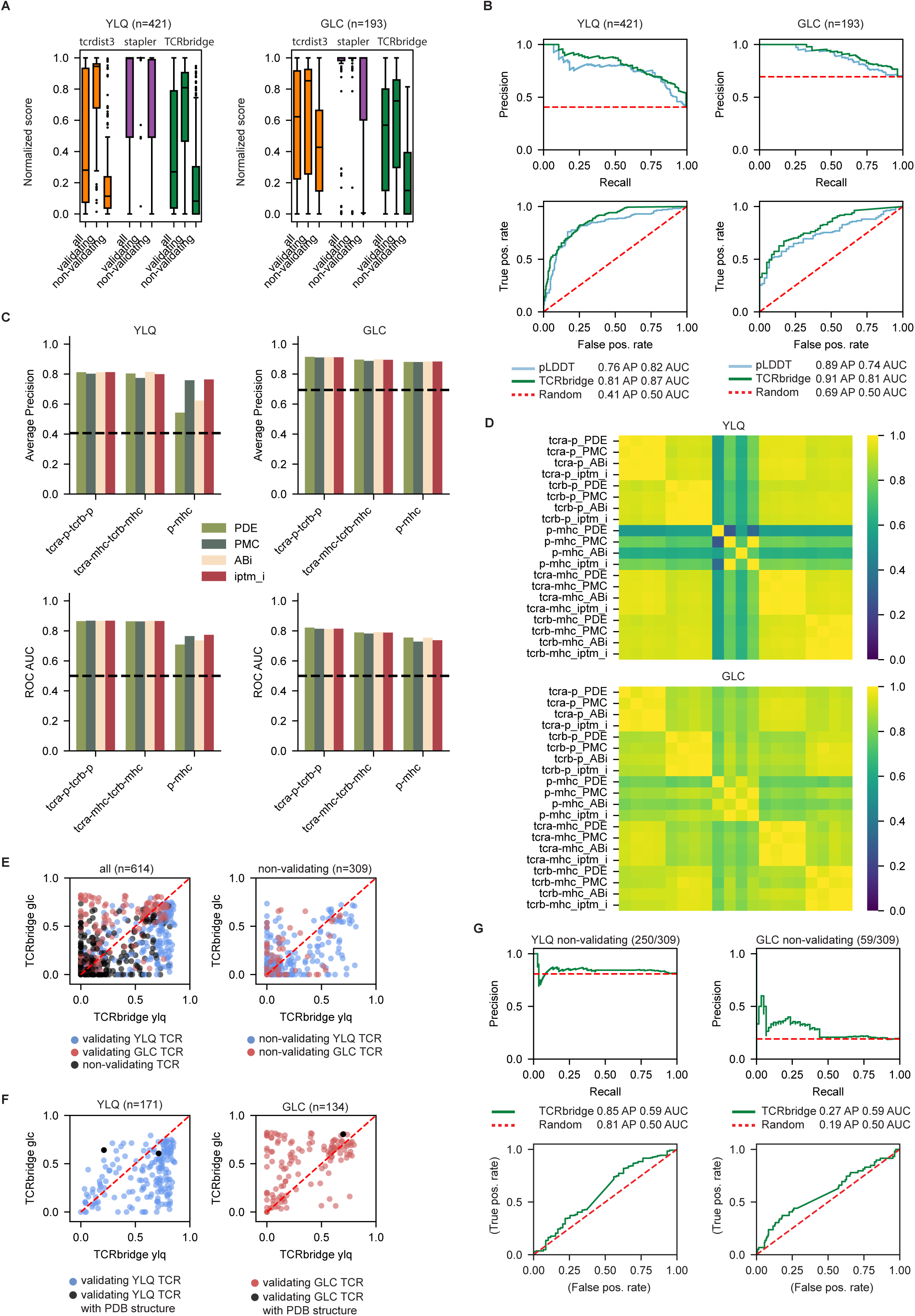
Analysis of model prediction scores. (A) Boxplot of tcrdist3, STAPLER, and TCRbridge normalized predicted confidence in TCRs being reactive. Confidence scores were normalized by dividing by the maximal prediction score for each model. Boxes indicate model median and 25th/75th percentiles of the normalized scores, whiskers show min/max except for outliers determined using the inter quartile range (IQR). (B) Comparison of the performance of TCRbridge and AlphaFold3 (AF3) mean pLDDT (predicted Local Distance Difference Test) confidence in the CDR3αβ and peptide regions of predicted TCR-pMHC complex structure in distinguishing validating and non-validating YLQ and GLC TCRs. Interpretation of Precision-Recall and ROC curves is described in Figure 4D. (C) Comparison of the performance of the indicated AF3 per-residue confidence scores and their combinations in predicted interfaces between the TCR and pMHC in distinguishing validating and non-validating YLQ and GLC TCRs. Performance of the following AF3 confidence metrics in its predicted interface were analyzed: interface predicted TM-score (ipTM), predicted distance error (PDE), and predicted merged confidence (PMC)—a combination of predicted aligned error (PAE) and pLDDT as performed by AlphaBridge^31^. TCRbridge uses ABi, which is formed by the combination of PDE and PMC scores. We evaluated performance using the sum of confidence in the TCRα-peptide and TCRβ-peptide interfaces (tcra-p-tcrb-p), the sum of confidence in the TCRα-MHC and TCRβ-MHC interfaces (tcra-mhc-tcrb-mhc), or the confidence in the peptide – MHC (p-mhc) interface. (D) Spearman correlation of confidence scores for the indicated AF3 and AlphaBridge metrics for predicted TCR-pMHC interfaces. Used confidence metrics and interfaces are described in (C). (E) Scatterplots depicting TCRbridge-predicted confidence scores in TCR-pMHC interfaces of all YLQ– and GLC-annotated (i.e., validating and non-validating, left) and of non-validating YLQ and GLC (right) TCRs in predicted structures with either their cognate or non-cognate epitopes. (F) Scatterplots depicting TCRbridge-predicted confidence scores in TCR-pMHC interfaces of validating YLQ (left) and validating GLC (right) TCRs in predicted structures with either their cognate or non-cognate epitopes. Dots representing TCRs deposited in PDB before the AlphaFold3 training cutoff date are highlighted in black. (G) Performance of TCRbridge in scoring predicted structures with presumed cognate peptides and non-cognate peptides for non-validating TCRs. Precision-recall and ROC curves show performance for non-validating YLQ (top) and GLC (bottom) TCRs. Note the limited performance for non-validating TCRs as compared to that observed for validating TCRs, as described in Figure 4E. Dashed red lines represent random performance.

## Supplemental Information

Table S1 Parsed VDJdb database

Table S2 Well-specific orthogonal primers

Table S3 Plate-specific orthogonal primers

Table S4 TRAV and TRBV sequences

Table S5 Nanopore primers

